# Unravelling the metastasis-preventing effect of miR-200c *in vitro* and *in vivo*

**DOI:** 10.1101/2023.11.14.566527

**Authors:** Bianca Köhler, Emily Brieger, Tom Brandstätter, Elisa Hörterer, Ulrich Wilk, Jana Pöhmerer, Anna Jötten, Philipp Paulitschke, Chase P Broedersz, Stefan Zahler, Joachim O Rädler, Ernst Wagner, Andreas Roidl

## Abstract

Advanced breast cancer as well as insufficient treatment can lead to the dissemination of malignant cells from the primary tumor to distant organs. Recent research has shown that miR-200c can hamper certain steps of the invasion-metastasis cascade. However, it is still unclear, whether sole miR-200c expression is sufficient to prevent breast cancer cells from metastasis formation. Hence, we performed a xenograft mouse experiment with inducible miR-200c expression in MDA-MB 231 cells. The *ex vivo* analysis of metastatic sites in a multitude of organs including lung, liver, brain, and spleen has revealed a dramatically reduced metastatic burden of mice with miR-200c expressing tumors. A fundamental prerequisite for metastasis formation is the motility of cancer cells and, therefore, their migration. Consequently, we analyzed the effect of miR-200c on collective and single cell migration *in vitro*, utilizing MDA-MB 231 and MCF7 cell systems with genetically modified miR-200c expression. Analysis of collective cell migration has resulted in confluence dependent motility of cells with altered miR-200c expression. Additionally, scratch assays have shown enhanced predisposition of miR-200c negative cells to leave cell clusters. The in-between stage of collective and single cell migration was validated using transwell assays, which have displayed reduced migration of miR-200c positive cells. Finally, to measure migration on single cell level, a novel assay on dumbbell shaped micropatterns was performed, which revealed that miR-200c critically determines confined cell motility. All of these results demonstrate that exclusive expression of miR-200c impedes metastasis formation *in vivo* and migration *in vitro* and highlight miR-200c as metastatic suppressor in breast cancer.

## Introduction

Metastasis is the process of spreading of malignant cells from a primary tumor to distant organs (1, 2). It is the leading cause for about 90 % of cancer-related deaths (3, 4), and in regard to fundamental biological functions, insufficiently understood (5–7). In most cases (60 to 70 % of cancer patients), the metastatic process has already been initiated at the time of diagnosis (8). To prevent or even reverse metastasis in a targeted manner, it is therefore, essential to better elucidate this process (9).

Metastases are the consequences of a multi-step process, also known as the invasion-metastasis cascade (9). This cascade comprises of the following interrelated steps: invasion, intravasation, circulation, extravasation, and metastatic colonization of malignant cells (6, 8–11). One indispensable step of metastasis formation is the ability of cancer cells to migrate (12, 13). After initial invasion through the basement membrane, cancer cells migrate through different microenvironments i.e., tissue, stroma, blood vessels and lymphatics to disseminate into pre-metastatic niches in distant organs (11, 14, 15). To reach these tissues, cancer cells can either migrate individually or collectively (7, 12, 14, 16, 17). The most common type is collective migration where cells within the clusters communicate with each other and cell-cell adhesion is maintained (7, 11). In contrast to this, during single cell migration, a solitary cell detaches from the tumor to reach a distant site (10, 11).

Finding factors that modulate cell motility and thus metastasis is essential to reduce the metastatic burden of patients. One promising mechanism to prevent metastasis formation is the regulation of the expression of small non-coding RNAs, especially microRNAs (miRNAs). These RNAs alter the expression of target messenger RNAs through binding to the three prime untranslated region (3’-UTR). This results in either the degradation or the translational inhibition of the target mRNA (18, 19). MiRNAs have the potential to target multiple mRNAs at the same time (20) resulting in complex miRNA-mRNA-interactions. In previous studies, we and others extensively studied the role of the tumor suppressor microRNA 200c (miR-200c), which is involved in many processes of tumorigenesis, e.g. apoptosis, proliferation, chemotherapy resistance, motility, epithelial-mesenchymal transition (EMT), and metastasis (21–26). MiR-200c belongs to the microRNA-200 family. Its five members are located on two chromosomes: miR-200b, -200a, and -429 on chromosome 1p36.33 and miR-200c, and -141 on chromosome 12p13.31. Together with miR-200b, and -429, miR-200c possesses the same seed region (1). MicroRNA 200c is a very attractive therapeutic molecule as it can play a role in every step of the invasion-metastasis cascade, e.g. tissue invasion, migration, and survival in circulation (anoikis), as well as in EMT (1, 18, 27–29). Many of its targets like ZEB1, ZEB2, USP25, MSN and FN1 alter migration directly or re-organize the actin cytoskeleton (FHOD1, FLNA and PPM1F) and thus can indirectly affect migration and invasion (28). *In vivo* studies of miR-200c overexpression showed a reduced tumor growth and prolonged survival (21, 30, 31), whereas metastases formation was investigated by overexpressing the miR-200c/141 cluster (32) or only focused on the analysis of lung metastases in claudin-low breast cancer (33).

Given that miRNAs possess a multitude of targets, we investigated whether exclusive miR-200c expression is sufficient to hamper tumor cell motility, and thus, metastasis formation. Induction of miR-200c in the cells of the primary tumor revealed a dramatic reduction of the metastatic burden. As migration is the prerequisite for metastasis formation, we revealed by several novel and established assays that miR-200c is a crucial player lowering the migratory capacity in tumor cells by harnessing intrinsic motility factors on cell clusters but also on single cell level. MiR-200c can thus function as a potential biomarker and metastasis suppressor and hence facilitates decision-making on effective breast cancer treatment of patients. Moreover, a successful delivery of miR-200c to primary tumors might increase patients’ survival and reduce recurrence rates.

## Material and Methods

### 1. Reagents

Doxycycline hyclate (cat. no. D9891) was purchased from Sigma-Aldrich (St. Louis, MI, USA) and was resolved in sterile RNase/DNase-free water (Sigma-Aldrich, St. Louis, MI, USA, cat. no. 95284-1L). Hoechst 33342 (cat. no. H1399) was acquired from Thermofisher (Thermofisher, Germany).

### 2. Cell Culture

The MDA-MB 231 cell line was acquired from DSMZ (Braunschweig, Germany) and utilized to generate both doxycycline inducible cell lines: MDA-MB 231 Tripz 200c and MDA-MB 231 Tripz Ctrl. The generation of the cells is described in Ljepoja *et al.* (22). A second lentiviral transduction was carried out in order to generate Luciferase tagged cell lines to enable *ex vivo* Luciferase analysis of the animals’ organs. For this purpose, a third generation lentiviral system with the packaging plasmids pRSV-Rev (addgene plasmid #12253), pMDLg/pRRE (addgene plasmid #12251) and pCMV-VSV-G (addgene plasmid #8454) and the transfer plasmid pLenti CMV Puro LUC (w168-1) (addgene plasmid #17477) was transduced. The pRSV-Rev and pMDLg/pRRE plasmids were a gift from Didier Trono, pCMV-VSV-G was a gift from Bob Weinberg and the transfer plasmid was a gift from Eric Campeau & Paul Kaufman (34–36). Subsequently, cells were selected with puromycin dihydrochloride (Sigma-Aldrich, St. Louis, MI, USA, cat. no. P8833-10MG) for 48 hours. Parental and genetically modified MDA-MB 231 cells were cultured at 37 °C and 0 % CO2 in Leibovitz’s L-15 medium (Sigma-Aldrich) supplemented with 10 % fetal calf serum (FCS, Gibco, ThermoFisher Scientific, Hanover Park, IL, USA). In order to induce the Tripz-construct cells were treated with a final concentration of 5 µg/ml doxycycline hyclate (DOX). Depletion of the DOX was prevented by adding fresh DOX to the medium every 48 to 72 hours. MCF7 wildtype (wt) cells were acquired from Cell Line Service (Eppelheim, Germany). The TALENs KO was conducted in our lab as previously described (23). All MCF7 cell lines were cultured at 37 °C and 5 % CO2 in high glucose DMEM medium (Sigma-Aldrich) supplemented with 10 % FCS (Gibco). Cells were routinely tested for mycoplasm contamination.

### 3. *In vitro* confined cell motility analysis (1D dumbbells)

Single cell motility was monitored on 1D dumbbell micropatterns. For micropatterning the surface of the ibiTreat µ-dish (ibidi, Germany, cat. no. 81156) was passivated with a small drop of 0.01 % (w/v) PLL (Sigma-Aldrich, cat. no. P8920-100ML). After incubating the dish for 30 minutes with PLL at room temperature, it was rinsed with HEPES buffer (pH = 8.3, Thermo scientific, cat. no. J16924-AP) and subsequently 100 mg/ml of mPEG-SVA (LaysanBio, cat. no. NC0107576) diluted in 0.1 M HEPES were evenly distributed. The dish was further incubated at room temperature for at least 1 hour before rinsing with miliQ water. The passivated dish was then photopatterned using the PRIMO module (Alvéole, France) mounted on an automated inverted microscope (Nikon Eclipse Ti). After passivation, PLPP gel (Alvéole) was diluted in 99 % ethanol to distribute the gel evenly throughout the surface. The dumbbell shaped pattern was placed on top of the dish via the Leonardo software (Alvéole) and illuminated with UV-light with a dose of 15 mJ/mm^2^. Next, the dish is washed with milliQ water and rehydrated with PBS for 5 minutes followed by an incubation with 20 µg/ml of labelled Fibronectin-Alexa647 (Y-proteins cat. no. 663, Thermofisher, cat. no. A37573) for 15 minutes at room temperature. Once the dish was washed with PBS a total number of 10,000 cells, when appropriate 72 hours treated with DOX prior to seeding, were added and left to adhere for at least 4 hours. After that, the medium was exchanged to medium without phenol red containing Hoechst 33342 at a final concentration of 25 nM for nuclear staining. If necessary, medium was additionally supplemented with 5 µg/ml DOX. All confined motility measurements were performed in time-lapse mode for 48 hours on a Nikon Eclipse Ti microscope using a 10× objective. The experiments were carried out at cells’ specific cultivating conditions using a heated chamber (Okolab) at 37 °C. Images (brightfield and DAPI) are acquired every 10 minutes. Cell tracking of the nuclei was performed by using TrackPy (Python Version 3.10.5). 89 tracks of the MDA-MB 231 Tripz Ctrl +DOX, 85 tracks of the MDA-MB 231 Tripz 200c and 94 tracks of the MDA-MB 231 Tripz 200c +DOX were analyzed. Each single measurement was conducted for at least 48 hours, but track length varied between 8 hours (minimum) and 48 hours (maximum). To quantify the dynamics of confined cell migration, we analyzed the stay probability (former also termed survival probability (37)) *S*(*t*). This is the probability that a cell has not transitioned from one island to the other after a time interval of length *t*. The stay probability was calculated from the distribution of dwell times *p*(τ). The dwell time τ here is the time a cell spent on its own island before it transitioned to the other island. Transition events were detected in the cell trajectories, allowing the measurement of τ. The stay probability was then computed through:

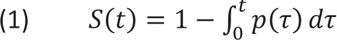

For more information, refer to Brückner *et al.* (37). Finally, from the trajectories, the speed of the cells on the bridge was measured through *v* = (*x*(*t* + Δ*t*) − *x*(*t*))/Δ*t*, where *x*(*t*) and *x*(*t* + Δ*t*) are two consecutive positions of the cell nucleus and Δ*t* is the measurement interval of tracking the cells. This results in an additional characterization of the hopping dynamics. The average of this speed was displayed, and error bars represent the error of the mean. To gain more detailed insight into the dynamics of cell hopping, a physical model for cell migration within our micropattern was inferred. The complexity of cell migration was reduced by employing a coarse-grained description of cell migration: we capture the dynamics for the nucleus position *x* and the nucleus velocity *v* using an underdamped Langevin equation, and used a previously developed method to directly infer such an equation from experimental data (37):

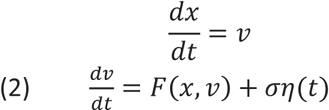

Here *F*(*x*, *v*) is an effective force that describes the deterministic component of how the cell nucleus is accelerated at a given position and velocity. This effective force encodes how the migrating cell interacts with the geometric confinement of the micropattern. To capture the stochasticity of cell migration, leading to a wide variety of different cell behaviors, a dynamical noise term ση(*t*) is included that adds to the acceleration of the nucleus, where σ is the noise amplitude and η(*t*) is a Gaussian white noise with ⟨η(*t*)⟩ = 0 and ⟨η(*t*)η(*t*′)⟩ = δ(*t* − *t*′). Using a statistical learning framework called Underdamped Langevin Inference (ULI) (38), the deterministic and the stochastic terms in the Langevin equation were inferred from our experimental trajectory data. This approach has the advantage that a specific form for *F*(*x*, *v*) and a specific value for σ were not a priori assumed based on physical principles. Instead, the experimental rigorously constrained the form and value of these terms. We refer to Brückner *et al.* for more details (38). From the inferred model, the structure of *F*(*x*, *v*) was analyzed, which provided information on the underlying qualitative features of the hopping dynamics as represented.

### 4. Transwell assays

Migration analysis was performed in a transwell assay using Falcon Cell Culture Inserts with a membrane pore size of 8 µm (Corning, NC, USA, cat. no. 353097) in a 24 well format. Inserts (n = 3) were filled with 500 µl of appropriate medium supplemented with 0.5 % FCS and 100,000 cells. MDA-MB 231 Tripz 200c cells were either 72 hours pre-induced with DOX (5 µg/ml) before seeding and induced with DOX as mentioned or not. Wells were filled with 750 µl corresponding medium supplemented with 10 % FCS. Cells were incubated 18 hours and subsequently excess of the cell suspension was removed and the insert was carefully washed with PBS. For permeabilization, inserts were incubated 20 minutes with 100 % methanol. After washing, inserts were placed into 0.5 % crystal violet solution containing 25 % methanol for 15 minutes. Excess of the solution was removed by washing the inserts several times with water and subsequently the inserts were dried overnight. Microscopic pictures of migrated cells were taken with a 5× magnification of the central part of the membrane. These pictures were analyzed using the ImageJ 1.53e software. To determine the quantity of migrated cells, pictures were binarized and the “area” of black pixels was calculated. Relative migration is presented by normalization to the wildtype cell line. Finally, transwell membranes were incubated in 96 % ethanol for 10 minutes. The dissolved crystal violet was subsequently measured with a TECAN reader at 595 nm (duplicates per insert). The mean of each experiment is displayed as a violin plot.

### 5. Random walk and scratch assays

For the random walk experiments cells were seeded with a concentration of 50,000 cells per well (n = 3). MDA-MB 231 Tripz 200c and Ctrl were pretreated with DOX for 72 hours before seeding if appropriate. Alterations in the motility of the cells were monitored using a PHIO Cellwatcher M (PHIO scientific GmbH, Munich, Germany). Motility behavior of the cells was evaluated between 30 and 80 % confluency. Motility in µm/h was described for this condition as well as the mean motility of the triplicates for every specific confluency. Additionally, pictures of the MCF7 wt and KO 200c clusters formed after 50 hours of cultivation were taken using the Cellwatcher M. Scratch assays were performed in 8-well coverslides (ibidi, Germany, cat. no. 80826) and cells were seeded 24 hours prior to performing the scratch in 100 % confluent cells. MDA-MB 231 Tripz 200c cells were pre-induced for 72 hours with DOX when appropriate. Nuclei of the cells were stained with the live cell imaging probe siR-DNA, Spirochrome (tebu-bio, Germany, cat. no. 251 SC007). Cells were stained with 1 µM siR-DNA solution and subsequently migratory behavior of the cells was recorded under a confocal-microscope (Leica SP8) for 24 hours. Microscope settings: A 10×/0.30 DRY objective and hybrid detector (Leica HyD 641 nm – 777 nm) for siR-DNA fluorescence detection and a photomultiplier (PMT) were applied. The Diode 638 laser for the laser line of 638 nm was used. Live cell imaging was conducted with a frame rate of 1 per 5 minutes. Data was obtained with the LAS X software 3.5.7.23225. Migration was quantified by determining the closure of the scratch. Therefore, occupied area by cells was defined at 0 and 10 hours after scratching for the MDA-MB 231 cells and 14 and 24 hours after scratching for the MCF7 cells. All cells were stained as mentioned previously and pictures show siR-DNA labeled cells at the indicated timepoints. The difference in area in % was quantified using the Fiji plug in “Wound healing size tool” (39). The blue lines define the boarders of the scratch which were automatically calculated with the same parameters within each live cell imaging sample. Additionally, the number of invaded cells as well as the confinement ratio were analyzed. To be able to identify invaded cells, blue lines were hand-drawn indicating the wound-boarder at 0 hour (MDA-MB 231) and 14 hours (MCF7) after scratch performance. The number of cells which invaded into the free area of the scratch was calculated manually after 10 hours. The confinement ratio of each cell was quantified by converting the images to 8bit, using bandpass filter and subtracting the background. Subsequently the Fiji plug in “TrackMate” (DOG detector 15 pix., LAP tracker 15 pix.) was utilized to obtain the tracks (40). Motility behavior of the MDA-MB 231 cells was further analyzed by dividing the cells within a scratch experiment in cells of the “migratory front” and the dense cell clusters named “bulk”. The migratory behavior of cells of these two regions (further subdivided into left and right side, n = 10) was quantified using the “Chemotaxis and Migration Tool” Version 2.0 from ibidi GmbH (Germany) by determining the directionality of the cells. To calculate the directionality of a cell the ratio of the Euclidian distance to the accumulated distance was used. Accumulated distance in µm was determined using the “Chemotaxis and Migration Tool” Version 2.0 (stand-alone) from ibidi GmbH (Germany). Position of the cells in X and Y axis was taken from the TrackMate analysis. Within each condition the movement of cells was summarized as the number of cells migrating to the right or left side (numbers indicated within the graphs). Red trajectories indicated movement of these cells into the scratch to close it. The difference between MDA-MB 231 cells with and without miR-200c expression was quantified by comparing the directionality of the front and bulk cells (n = 80 cells, 20 cells per biological replicate (4 replicates in total), right and left side were cumulated).

### 6. *In vivo* analysis of metastatic formation in distant organs of mice with or without miR-200c expressing breast cancer tumors

A total number of five million human MDA-MB 231 Tripz 200c Luc cells were injected s.c. into the left flank of twenty 6-week-old female NMRI-nu mice (Janvier, Le-Genest-St-Isle, France). Mice were randomized into two experimental groups when their tumor reached a size of approximately 200 mm^3^. One group was kept with the original, normal feed, whereas the second group of mice was switched to doxycycline containing feed (+ 625 mg/kg doxycycline, sniff Spezialdiäten, Soest, Germany, cat. no. A115 D70624) in order to induce miR-200c expression. Mice were kept with their corresponding diet until the end of the study. Additionally, a control study was performed to exclude DOX effects. Therefore, eighteen mice were randomized into two groups beforehand. One group was fed with normal feed (n = 7) and the second group of mice was nourished with DOX feed (n = 11) throughout the study. Five million MDA-MB 231 Tripz Ctrl Luc cells were injected into these mice. Tumor growth as well as animal well-being and weight were monitored over the entire period. Tumor growth was monitored throughout the whole study using caliper measurement and calculated with the following equation: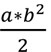 (a = longest side of the tumor; b = widest side vertical to a) (41). Criteria for euthanasia were set to a critical tumor size which was achieved when the tumor diameter was bigger than 12 mm. Mice were sacrificed by cervical dislocation when they reached the critical tumor size. If necessary, mice were sacrificed also before reaching the critical tumor size due to animal well-being reasons. All animal experiments were performed according to the guidelines of German law for the protection of animal life and were approved by the district government of Upper Bavaria. Reference number: ROB-55_2-2532_Vet_02-19-20. Immediately after euthanasia, mice were dissected, and their organs removed for *ex vivo* Luciferase analysis. Brain, lung, liver and spleen were further washed with PBS to remove any blood residuals and subsequently frozen overnight at -80 °C. Before luciferase activity measurement, organs were thawed and weight. Each organ was homogenized with a lysis solution containing a 1:5 dilution of Luciferase Cell Culture Lysis 5X Reagent (Promega, Madison, WI, USA, cat. no. E1531) in Millipore water supplemented with 1 % (v/v) protease and phosphatase inhibitor cocktail (Protease and Phosphatase Inhibitor Cocktail, Sigma-Aldrich, cat. no. PPC1010). Homogenization was carried out in different rounds: 4 times with 6 m/s and 2 times with 6.5 m/s intermitted by cooling on ice using MP Biomedicals™ Lysematrix D (ThermoFisher Scientific, Waltham, MA, USA, cat. no. 11432420) in a homogenizer (MP Biomedicals™ FastPrep-24, ThermoFisher Scientific, cat. no. 12079310). Samples were frozen overnight at -80 °C to allow further cell lysis. Test specimens were thawed and subsequently centrifuged at 13,300 rpm for 10 minutes at 4 °C. Luciferase activity was measured in 50 µl of the appropriate supernatant together with 100 µl of LAR buffer solution with 5 % (v/v) of a mixture of 10 mM luciferin and 29.375 mM glycylglycine. LAR buffer components: 20 mM glycylglycine, 1.0 mM MgCl2, 0.10 mM EDTA, 3.3 mM DTT, 0.55 mM ATP, and 0.27 mM coenzyme A adjusted to a pH of 8 – 8.5. Measurements were carried out in triplicates. Relative light units per gram organ (RLUs/g) were calculated as mean with SD. The livers were measured in two separate runs due to their greater weight and RLUs were finally calculated for the whole organ. Values smaller or equal to the values of the control measurement (solely lysis buffer) were set to zero and the organ was termed metastases free.

### 7. Software

The figures 1A and B as well as the figure 4G were created with BioRender.com (accessed on 7^th^ of November 2023).

**Figure 1:**
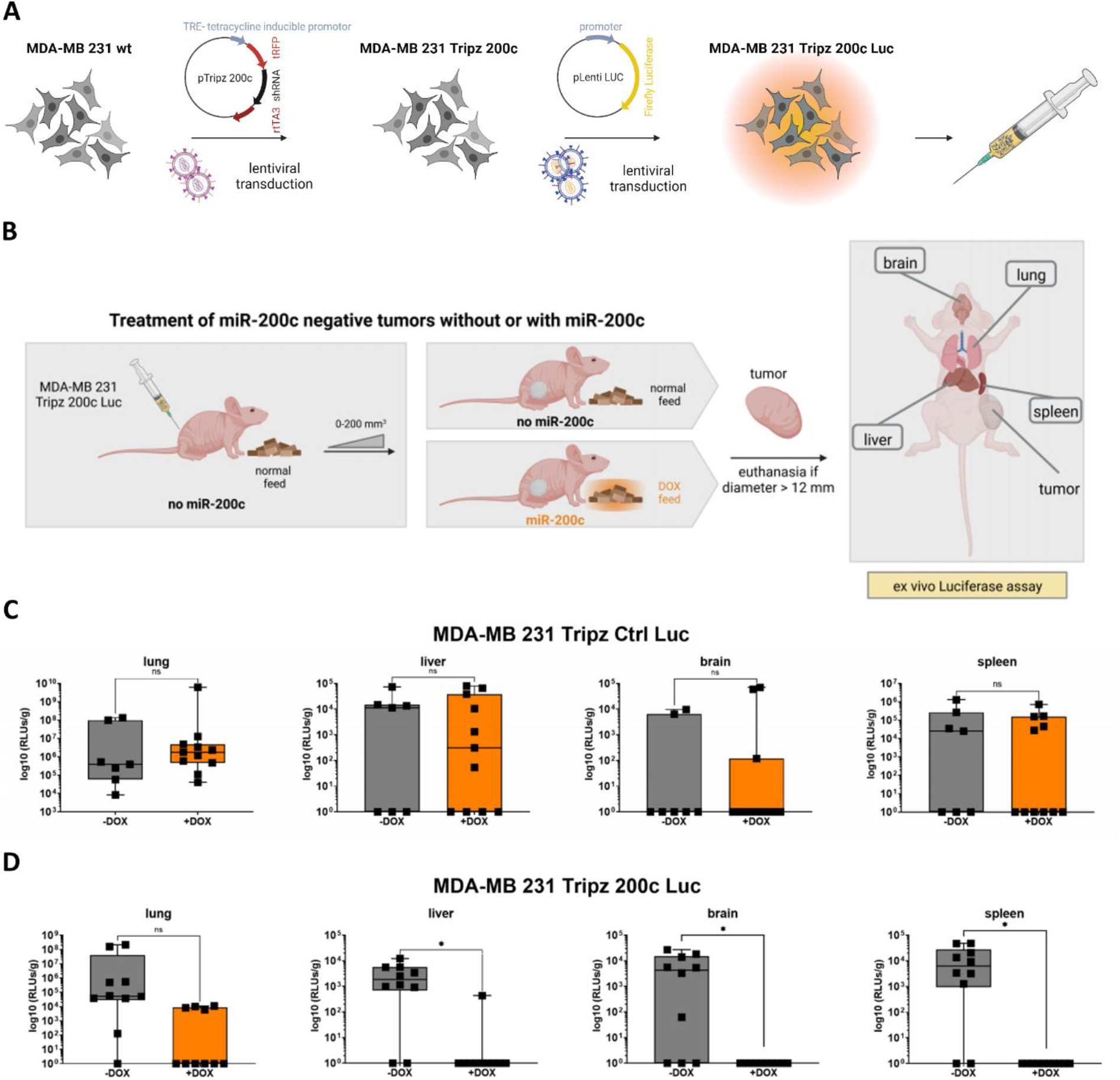
Metastasis formation in mice is reduced when miR-200c is expressed in the primary tumor. **(A)** Generation of the doxycycline inducible MDA-MB 231 Tripz 200c cell line which was further transduced with a luciferase tag, using a lentiviral system, for injection into mice. Figure was created with BioRender.com. **(B)** Experimental procedure for the treatment of inoculated MDA-MB 231 Tripz 200c Luc cells in mice. After initial tumor growth with normal feed till approximately a size of 200 mm^3^ mice were randomized into two diet groups: doxycycline (DOX) containing or further normal feed. Mice were euthanized as indicated and different organs were analyzed for metastasis formation performing an *ex vivo* luciferase assay. Figure was created with BioRender.com. **(C)** Quantification of luminescence (relative light units, RLUs) and therefore metastasis formation in lung, liver, brain and spleen of mice of a control group. MDA-MB 231 Tripz Ctrl Luc cells were injected into the mice of this control group. These breast cancer cells express a scrambled control sequence upon doxycycline induction. Cell generation is in accordance with the previously described procedure. Mice were either fed with normal (gray, n = 7) or doxycycline feed (orange, n = 11) ab initio. Values are displayed on a logarithmic scale as relative light units (RLUs) per gram organ (RLUs/g). Each data point represents one mouse. For statistical evaluation an unpaired, two tailed student’s t-test was performed. ns = not significant. **(D)** Quantification of luminescence (relative light units, RLUs) and therefore metastasis formation in lung, liver, brain and spleen of mice of MDA-MB 231 Tripz 200c Luc cells without (gray, n = 10) or with miR-200c expression (orange, n = 10). Change of diet was conducted as described previously. Values are displayed on a logarithmic scale as relative light units (RLUs) per gram organ (RLUs/g). Each data point represents one mouse. For statistical evaluation an unpaired, two tailed student’s t-test was performed. ns = not significant, * p < 0.05.

### 8. Statistical Analysis

Statistical significance was calculated utilizing GraphPad Prism 7.04. To compare two samples an unpaired two-tailed Student’s t-test was used. For comparing more than two samples a two-way ANOVA statistical test was conducted. * p < 0.05, ** p < 0.01, *** p < 0.001, **** p < 0.0001. Values are displayed as mean with SD.

## Results

### 1. MiR-200c lowers the metastatic burden *in vivo*

To investigate whether the exclusive expression of miR-200c is sufficient to prevent metastases, we utilized the triple negative breast cancer (TNBC) cell line MDA-MB 231, which was genetically modified with a Tripz-construct to selectively induce miR-200c expression upon treatment with doxycycline (DOX) (22). In order to monitor disseminated cells from the primary tumor this cell line was additionally transduced with a luciferase tag (Figure 1A). These cells were injected into mice and after initial tumor growth to a size of 200 mm^3^, mice were randomized into two diet groups (n = 10) (Figure 1B). Mice were either kept with their normal diet, where the inducible construct was not activated, or fed with DOX-containing diet to express miR-200c in the inoculated cells. We analyzed the metastatic spread of the primary tumor of each mouse *ex vivo* in the distant organs lung, liver, brain and spleen. Except for the spleen, these organs are predominant sites for metastasis formation of primary breast cancer in patients, also known as “organotropic metastasis” (42, 43). Additionally, we conducted a xenograft mouse experiment with a control cell line (MDA-MB 231 Tripz Ctrl Luc), which expresses a scrambled sequence after DOX administration. We performed this approach in order to exclude DOX effects on metastasis formation. Here, regardless of the treatment group, all control mice showed metastases in the lung, liver, brain and spleen to a similar extent (Figure 1C) and therefore no effect of the doxycycline administration was detectable. Subsequently, we measured luciferase activity in the organs of mice with the inducible MDA-MB 231 Tripz 200c Luc primary tumors (Figure 1D). While in 90 % of the animals with tumors lacking miR-200c expression (gray) metastases were found in the lungs, only 40 % of the mice with miR-200c expressing tumors (orange) developed pulmonary metastases. Moreover, we found statistically significant differences in the metastatic spread for all remaining organs. 80 % of the mice without miR-200c expression showed metastases in the liver while only 10 % of miR-200c positive tumors spread into this organ. MiR-200c negative primary tumors formed metastases in 70 % of the cases in the brain and in 8 out of 10 mice in the spleen. Of note, all mice with miR-200c positive tumors were free of metastases in these two organs. Additionally, we evaluated the mean-difference in survival time, tumor volume and metastatic burden in mice bearing miR-200c depleted or expressing primary tumors. While we found no substantial difference in the control mice (with or without DOX diet, Figure 2A), considerable difference was detected when comparing the two groups from the miR-200c xenograft mouse model. The mean survival time of mice with miR-200c positive primary tumors was almost doubled compared to mice with miR-200c non-expressing tumors (Figure 2B left). Despite the increased survival time of miR-200c positive tumor bearing mice the tumor volume at day of euthanasia is significantly decreased (Figure 2B middle). Finally, the metastatic burden defined by the number of organs affected by metastases was analyzed. MiR-200c positive tumors spread on average into one distant organ whereas miR-200c non-expressing tumors affected three organs (Figure 2B right). To determine whether the elevated survival time or tumor size influences the metastasis capacity, we evaluated and compared these parameters for each individual mouse (Figure 2C to F). Conducting this analysis with the control mice no clear correlations were detected (Figure 2C and D). Both control groups (MDA-MB 231 Tripz Ctrl -DOX and +DOX) show comparable survival times and the corresponding metastatic burdens are evenly distributed (Figure 2C). The same applies to the comparison of tumor volume and metastatic burden in these two groups (Figure 2D). In contrast, while the survival time of mice with miR-200c positive tumors was on average twice as high as of miR-200c negative animals, the metastatic burden showed an inverse correlation (Figure 2E). This means, although miR-200c positive animals lived longer and thus there was more time for tumors to metastasize, many animals did not show any metastases (the metastatic burden mostly equaled zero). Comparing the tumor volume of miR-200c negative or positive tumors in relation to the metastatic burden, smaller tumors (miR-200c positive tumors) generally tend to show a lower metastatic burden (Figure 2F). However, by comparing similar tumor sizes of miR-200c negative and positive tumors, only miR-200c expressing ones displayed a low metastatic burden. In summary, the xenograft experiment clearly showed that miR-200c reduces the metastatic burden and thus is able to prevent metastasis formation *in vivo*.

**Figure 2:**
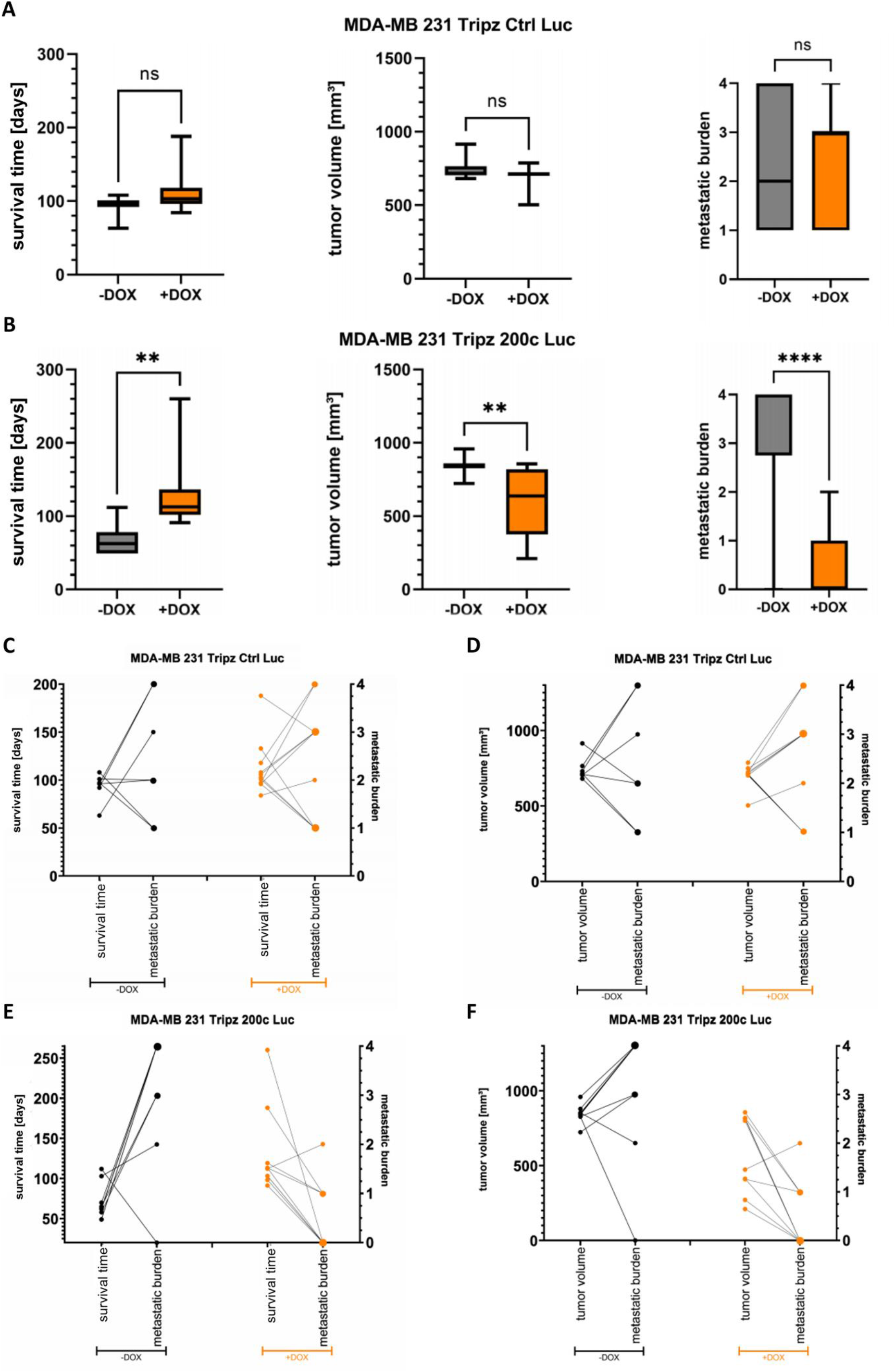
MiR-200c positively alters survival time of mice whereas it negatively influences tumor volume and metastatic burden. The term “metastatic burden” refers to the number of affected organs. **(A)** Survival time, tumor volume and metastatic burden of control mice injected with MDA-MB 231 Tripz Ctrl Luc cells. Mice were either fed with normal feed (gray, n = 7) or with doxycycline containing feed (orange, n = 11) ab initio. For statistical evaluation an unpaired, two tailed student’s t-test was performed. ns = not significant. **(B)** Survival time, tumor volume and metastatic burden of mice injected with MDA-MB 231 Tripz 200c Luc cells. Mice were either fed with normal feed (gray, n = 10) or with doxycycline containing feed (orange, miR-200c positive, n = 10) as indicated previously. For statistical evaluation an unpaired, two tailed student’s t-test was performed. ** p < 0.01 and **** p < 0.0001. Presentation of **(C)** survival time compared to the metastatic burden and **(D)** tumor volume compared to the metastatic burden of every individual control mouse. Results are separated into normal (black dots, n = 7) or doxycycline containing feed (orange dots, n = 11) group. Evaluation of **(E)** survival time compared to the metastatic burden and **(F)** tumor volume compared to the metastatic burden of every individual mouse with (orange) or without (gray) miR-200c expression of the primary tumor. Results are separated into normal (black dots, n = 10) or doxycycline containing feed (orange dots, n = 10) group. The bigger the dots the more frequently the same value of the metastatic burden was achieved.

### 2. MiR-200c expression modulates undirected collective migration

A prerequisite for metastasis formation is the motility of cells and thus cellular migration. One can discriminate between single cell migration and collective cell migration, where the movement of a single cell in a cell cluster is influenced by factors like cellular surface adhesion, cell-cell contacts or collisions, and gradients of chemoattractants. Our first *in vitro* assay sought to determine the impact of miR-200c on the random walk of the triple negative breast cancer cell line (MDA-MB 231) and the Luminal A breast cancer cells (MCF7) growing under regular culture conditions. While cells at low confluence migrate nearly unaffected by other cells, similar to single cell migration, higher confluence leads to cluster formation of cells resulting in collective cell migration behavior. By live imaging we were able to monitor cell migration during different confluence rates. A significant difference in the average motility was measured in the epithelial Luminal A breast cancer cell lines MCF7 wt and MCF7 KO 200c (Figure 3A). We observed MCF7 KO 200c cells having a higher motility rate when the confluence was lower (Figure 3B, Video 1 and 2), but this motility rate decreased when these cells became denser and finally approximated the motility rate of the MCF7 wt cells. Other than MCF7 wt, the mean motility of the MCF7 KO 200c cells seems to be dependent on the confluence. Moreover, pictures of the naturally formed cell clusters show big clusters in MCF7 wt cells while MCF7 KO 200c cells have smaller clusters and display more frequently single cells (Figure 3C). In contrast, the mesenchymal triple negative breast cancer cell line MDA-MB 231 Tripz 200c shows an opposite effect in its migratory behavior. The overall motility rate is slightly increased when these cells express miR-200c (Tripz +DOX) (Figure 3D, Video 3 and 4). When correlating motility rates with the confluence over time, miR-200c positive cells show increased motility compared to miR-200c negative cells with increasing cell density (Figure 3E). Overall, MDA-MB 231 cells show a higher motility rate compared to MCF7 cells and display an elevated motility rate the denser the cells grow (Figure 3E). These observed effects are not DOX induced as in MDA-MB 231 Tripz Ctrl cells no differences in the motility behavior of these two states were detected (Figure 3F and G). Altogether, in these experiments we observed that miR-200c positive cells in both cell systems behaved differently. Thus, it is very likely that the collective migration is influenced by factors as cell-cell-contacts, adhesion or the epithelial or mesenchymal phenotype, all parameters potentially affected by miR-200c.

**Figure 3:**
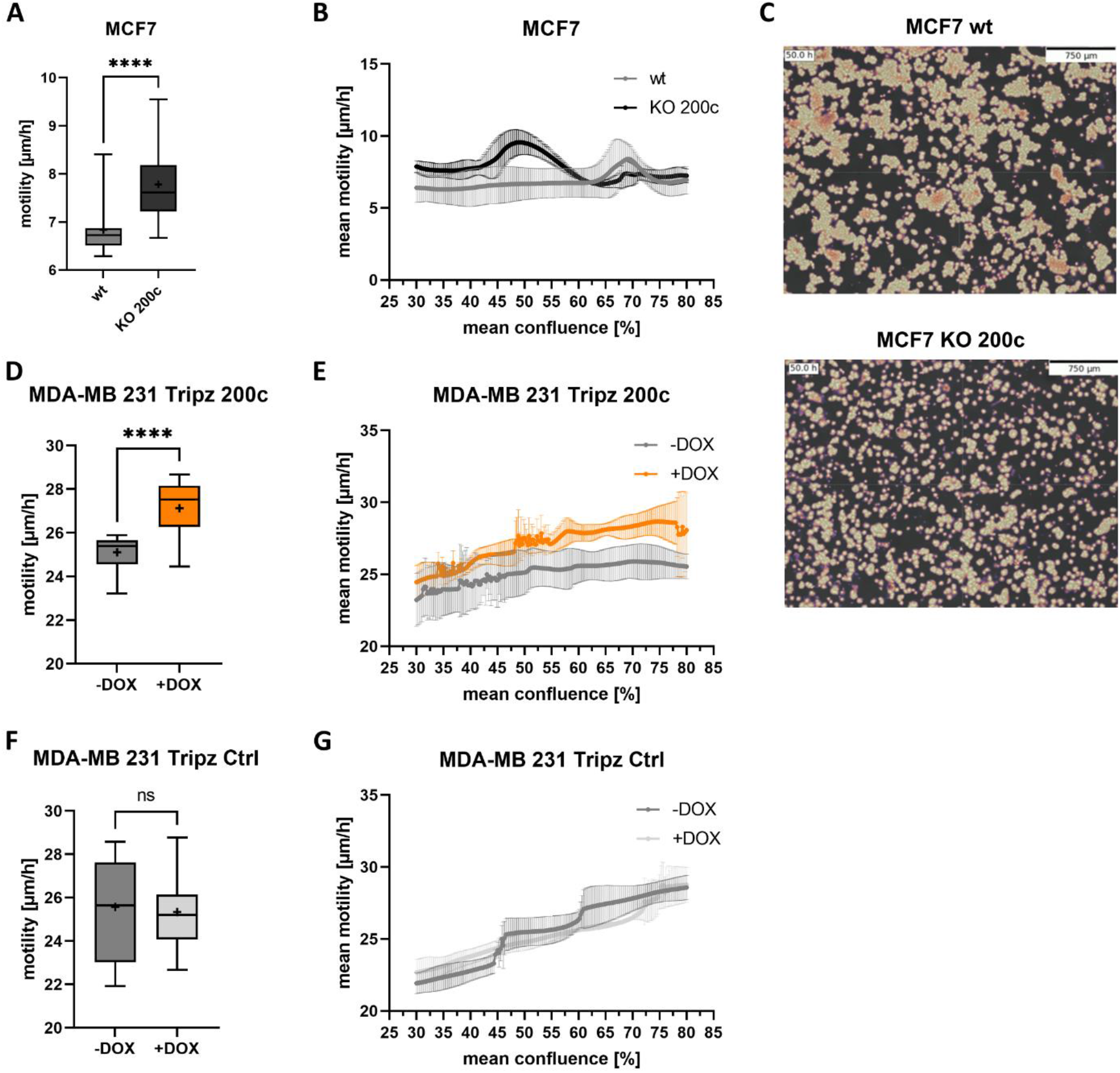
Mean motility is modulated by miR-200c expression dependent on the cellular phenotype. Motility behavior in random walk experiments of epithelial MCF7 wt and KO 200c cells. **(A)** Average motility in µm per hour [µm/h] (n = 3) and **(B)** mean motility [µm/h] at a specific confluence ranging from 30 to 80 % (n = 3). **(C)** Formed cell clusters of MCF7 wt (top) and MCF7 KO 200c cell (bottom) after 50 hours post seeding for the random walk analysis. Scale bar equals 750 µm. Pictures were taken with a Cellwatcher M device. **(D)** Average motility of MDA-MB 231 Tripz 200c cells with or without DOX induction in [µm/h] (n = 3) and **(E)** mean motility [µm/h] over a confluence rage of 30 to 80 % (n = 3). Control experiments to exclude any DOX effect was performed using MDA-MB 231 Tripz Ctrl cells. **(F)** Average motility of MDA-MB 231 Tripz Ctrl cells with or without DOX induction in [µm/h] (n = 3) and **(G)** mean motility [µm/h] over the confluence rage of 30 to 80 % (n = 3). Data of the average motility are presented as Box and whisker plots with minimal to maximal values. The median is plotted with a line and the mean with “+”. For statistical evaluation an unpaired, two tailed student’s t-test was performed. ns = not significant, **** p < 0.0001

**Figure 4:**
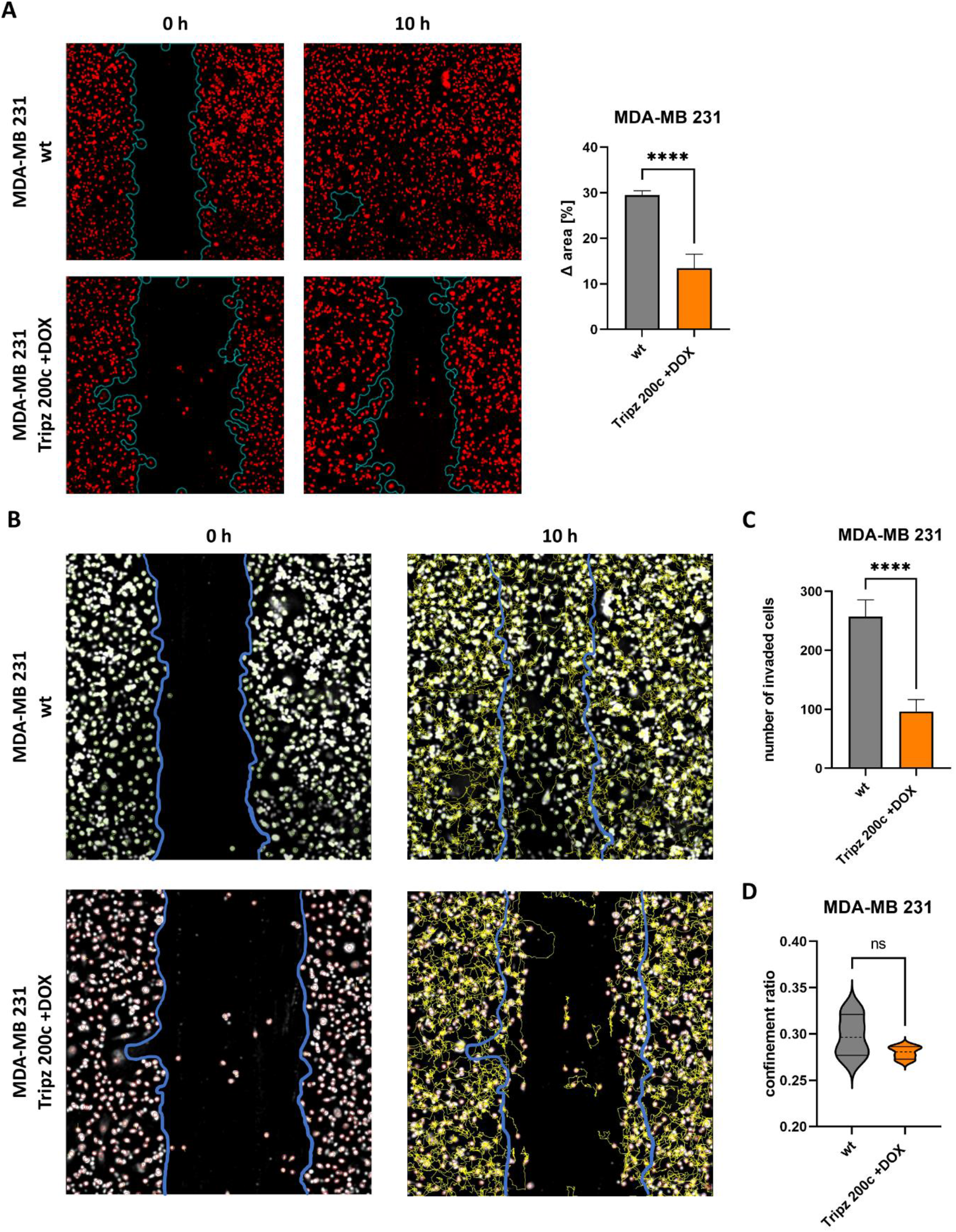
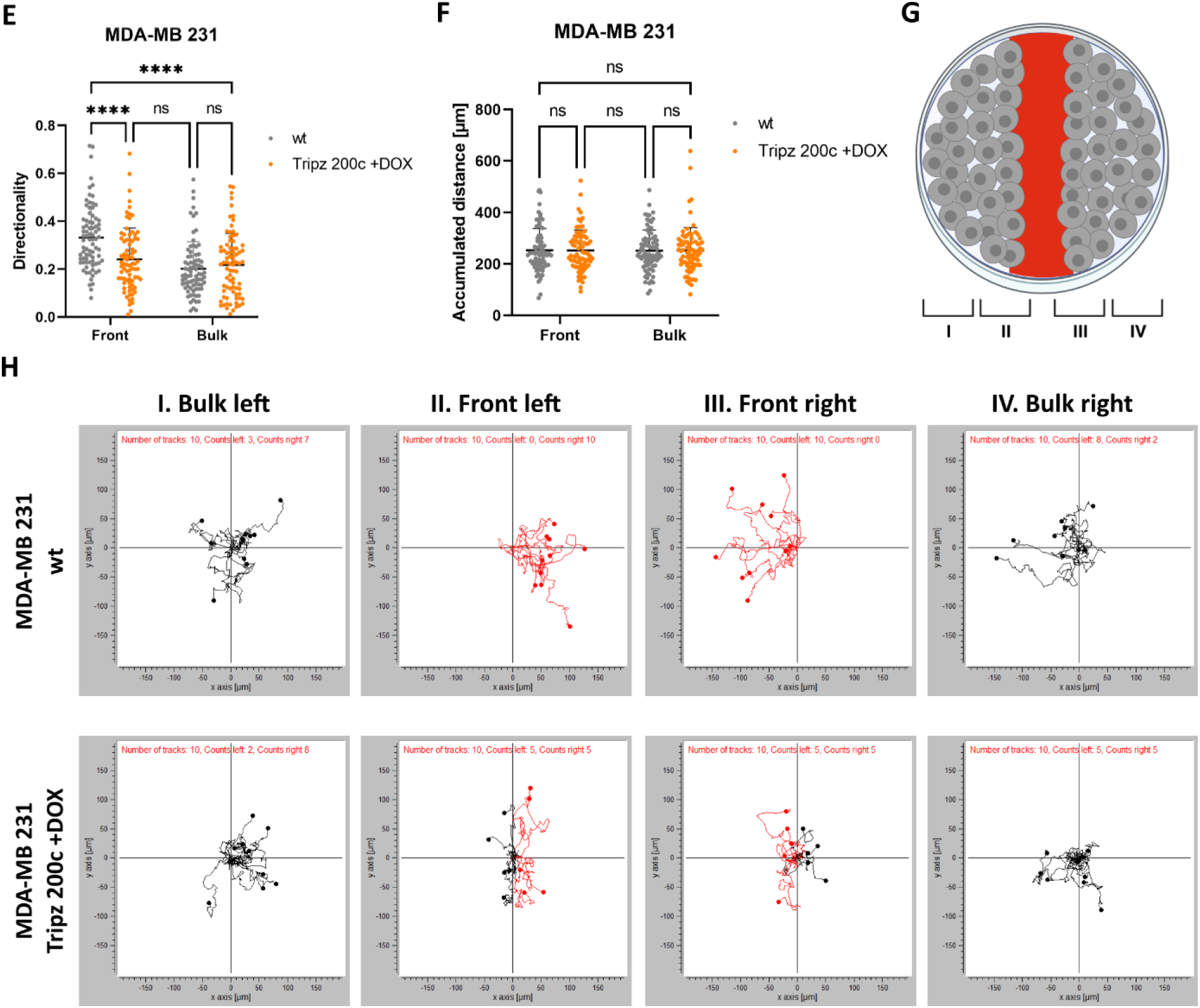
MDA-MB 231 breast cancer cells without miR-200c expression leave cell clusters more frequently. **(A)** Wound closure of MDA-MB 231 cell with (Tripz 200c +DOX) or without (wt) miR-200c expression. Blue lines in the microscopic pictures (left) indicate the borders of the wound at 0- and 10-hours post scratching. The difference in the area in [%] (right) is used to quantify the scratch closure (n = 4). Cells are stained with siR-DNA. **(B)** Microscopic pictures of MDA-MB 231 cells with different miR-200c expression for the analysis of **(C)** the number of invaded cells and **(D)** the confinement ratio (n = 4). Blue lines present the border of the scratch at 0 hours. Yellow lines represent the trajectories of the cells after 10 hours. For statistical evaluation an unpaired, two-tailed student’s t-test was performed. ns = not significant, **** p < 0.0001. The dashed line shows the median and the full lines the quartiles of the violin plot in (D). **(E)** Quantification of the directionality of MDA-MB 231 wt and induced Tripz 200c (+DOX) cells in the front and in the bulk region. For statistical evaluation a two-way ANOVA with multiple comparison and Šidák correction was performed. ns = not significant, ** p < 0.01, *** p < 0.001, **** p < 0.0001. **(F)** Quantification of the accumulated distance of MDA-MB 231 cells with altered miR-200c expression. For statistical evaluation a two-way ANOVA with multiple comparison and Šidák correction was performed. ns = not significant. (**G**) Analysis of migration behavior of cells in different regions of the scratch (front or bulk cells). Schematic presentation of the different areas within a scratch. Figure was created with BioRender.com. (**H**) Motility plots of cells in the bulk region left (I), front left (II), front right (III) and bulk right (IV) using the “Chemotaxis und Migration Tool” software from ibidi GmbH (Germany). Red trajectories represent migratory behavior into the scratch.

### 3. The migratory behavior in directed collective migration assays revealed enhanced predisposition of miR-200c non-expressing cells to leave cell clusters

To analyze collective cell migration in more detail, we performed scratch assays and measured the migration of cells from dense clusters into the free space. Wound closure, which is the increase of the occupied area within 10 hours, was significantly hampered in miR-200c induced MDA-MB 231 cells (Figure 4A, Video 5 to 8). Accordingly, MCF7 KO 200c cells inclined to close the wound faster than miR-200c expressing MCF7 wt cells (Supplementary Figure S1A, Supplementary Video S1 to S4). In both cellular systems significantly more cells invaded into the scratch when miR-200c was not expressed (Figure 4B and C and Supplementary Figure S1B and C). Additionally, the confinement ratio of miR-200c expressing and non-expressing cells was determined (Figure 4D and Supplementary Figure S1D). This ratio is defined as the cell’s net-distance divided by the total-distance traveled and represents the straightness of cell tracks (44, 45). In both cell lines overall no significant difference was measured (Figure 4D and Supplementary Figure S1D). However, MDA-MB 231 cells endogenously lacking miR-200c expression showed a two-parted distribution of the confinement ratio. This indicates a possible bilateral motility behavior, which might depend on the location of the cells within the scratch assay. For this reason, we performed a detailed evaluation of the cellular directionality at the front and the bulk regions (Figure 4E to H). We validated the trajectories of the MDA-MB 231 cells (Video 6 and 8) and observed an enhanced directionality of the wt MDA-MB 231 cells at the front regions compared to miR-200c expressing cells, bulk cells independent of their miR-200c expression level showed similar low directionality (Figure 4E). At the same time no significant difference in the accumulated distance was measured (Figure 4F). As an example, the movement of 10 cells per region I - IV (Figure 4G) visualizes the difference in how cells are closing the scratch (Figure 4H). In the bulk region of both miR-200c expression states no directed movement is observed. However, in the wt cell line 100 % of measured cells at both fronts are moving into the scratch, whereas only 50 % of miR-200c positive cells are contributing to the wound closure (Figure 4F). In summary, our data show an elevated intrinsic potential of cells without miR-200c to leave cell clusters making them more prone to disseminate into distant tissues.

### 4. Migration is reduced in miR-200c positive cells in transwell assays

Metastasis formation involves single cells disseminating from a cluster of cells, e.g. the primary tumor. Analogously, in a transwell assay a cell needs to disassociate from the cellular cluster and subsequently squeeze through a pore of a membrane. Therefore, this assay analyzes migration of both stages: collective cell migration and single cell migration. In order to pass through the pores cells are forced to alter their plasticity. The motility ability of cells in the transwell assay was determined by quantifying the number of cells which migrated through the pores of the membrane. The epithelial MCF7 wildtype cells migrated less frequently compared to the MCF7 KO 200c cells (Figure 5A). The quantitative analysis shows that the relative migration of MCF7 cells significantly increased by 1.5-fold when these cells did no longer express miR-200c (MCF7 KO 200c) (Figure 5B). Furthermore, the second cell system MDA-MB 231 Tripz 200c (-DOX) showed a higher migratory capacity, which was inhibited by 40 % when switching on miR-200c expression (+DOX) (Figure 5C and D). Both analyzes were additionally validated by determining the number of migrated cells by crystal violet staining (Supplementary Figure S2A and B). These data show, that besides an miR-200c-effect on collective migration, miR-200c acts negatively on single cell plasticity and motility.

**Figure 5:**
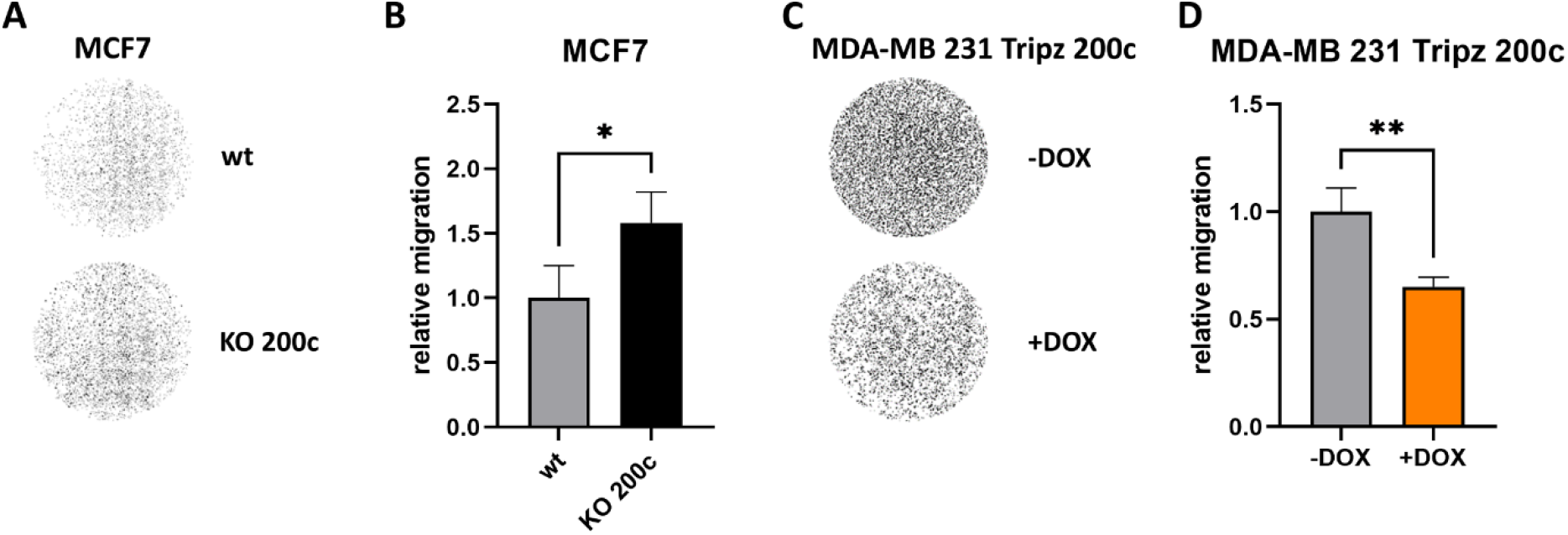
MiR-200c expression negatively modulates relative migration in a transwell assay. Pictures of the membranes with crystal violet stained, migrated **(A)** MCF7 wt and KO 200c cells and **(B)** the calculated relative migration (normalized to MCF7 wt cells, n = 3). With crystal violet stained **(C)** MDA-MB 231 Tripz 200c cells with and without DOX induction and **(D)** the corresponding quantification of the relative migration (normalized to MDA-MB 231 Tripz -DOX, n = 3). Values are displayed as mean with SD. For statistical evaluation an unpaired, two-tailed student’s t-test was performed. * p < 0.05, ** p < 0.01.

### 5. MiR-200c critically determines confined cell motility

To study a potential impact of miR-200c on single cell migration, we used a micropattern assay, consisting of two square islands connected by a thin bridge (Figure 6A and B). On this dumbbell-shaped assay, single cells repeatedly have to overcome the constricting bridge while they migrate. Therefore, with this assay, we gain quantitative insight into confined single cell migration, which is a key aspect of metastasis. On the micropattern, MDA-MB 231 cells were highly motile and repeatedly hopped between the two islands (Figure 6A and B, Video 9 and 10). Because this process is variable between repeated experiments (46), we imaged a large amount of micropatterns simultaneously (n> 85). To quantitatively study this inherently stochastic migration behavior, we tracked the cell nuclei in all micropatterns (Figure 6C and D). Using these trajectories, we analyzed the hopping behavior of both miR-200c negative (untreated, -DOX) and positive (treated with doxycycline, +DOX) MDA-MB 231 Tripz 200c cells. To quantify transition dynamics between the islands, we computed the stay probability *S(t)* that a cell was not performing a transition between islands after a certain time *t*. Upon induction of the miR-200c (+DOX), cells showed an approximately two-times higher probability not to make a transition (Figure 6E). This indicates less frequent transitions after induction of miR-200c, which correlates with the significant decrease in relative migration in our transwell assay (Figure 5D). Furthermore, we quantified the average speed of a cell when it is making a transition on the bridge. We found an approximately two-fold decrease of the transition speeds upon induction of miR-200c (Figure 6F). Taken together, these results showed that both transition frequency and transition speed of migrating MDA-MB-231 cells decreased due to miR-200c expression, indicating that miR-200c negatively affects the efficiency of confined cell migration.

**Figure 6:**
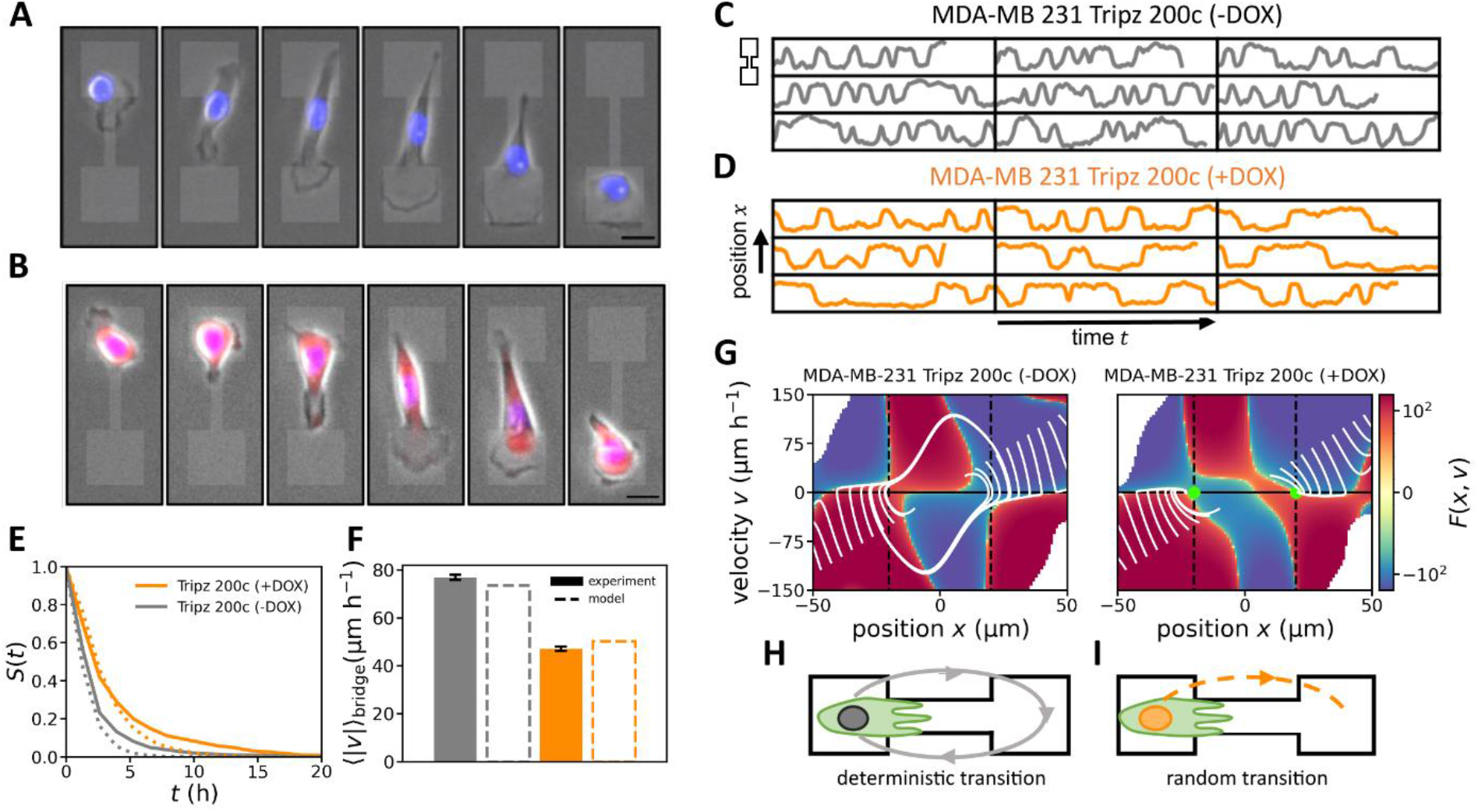
MiR-200c negatively affects the efficiency of confined cell migration. A) Time series of brightfield images of a MDA-MB 231 Tripz 200c cell without miR-200c expression performing hopping behavior in a two-state micropattern. **(B)** Time series of brightfield images of a MDA-MB 231 Tripz 200c cell with miR-200c expression. Nuclei are stained in blue. Scalebar equals 20 µm and applies to both (A) and (B). Red fluorescence shows the red fluorescence protein (RFP) which is simultaneously expressed with miR-200c upon effective DOX induction. Selection of nucleus trajectories for **(C)** MDA-MB 231 Tripz 200c (-DOX) and **(D)** MDA-MB 231 Tripz 200c (+DOX) cells. **(E)** Stay probability of the two cell lines which describes how likely it is that a cell has not made a transition after a certain time *t*. Both in (E) and (F), dotted lines indicate model results. **(F)** Average speed of the MDA-MB 231 Tripz 200c cells while making a transition on the bridge. Legend of (E) also applies here. **(G)** Inferred effective force *F*(*x*, *v*) describing the deterministic part of the hopping dynamics of the two states of the cells. White lines show the deterministic behavior in the two-dimensional phase space of the cells for the MDA-MB 231 cell line in both states. Green dots indicate the fix points of the dynamics in the (+DOX) case. Schematics in **(H)** and **(I)** show the qualitatively different dynamics of cells in the micropattern. Gray loop indicates deterministic transition dynamics where cells without miR-200c repeatedly oscillate between two islands. Dotted line indicates that cells with miR-200c only perform transitions into the opposite island when they are randomly excited to do so.

To further obtain a quantitative understanding of the hopping behavior, we aimed to describe cells in our micropattern using a dynamical model. We made use of a previously developed approach to directly infer from finite experimental trajectory data a Langevin equation of motion describing the stochastic nucleus trajectories (38). The key advantage of this inference procedure is that it can separate stochastic features from deterministic features of cell behavior. Specifically, we find an effective force *F*(*x*, *v*) for MDA-MB 231 cells with and without miR-200c expression. This effective force describes the deterministic part of the acceleration of the nucleus of these cells due to the average active cell migration behavior in our micropattern. The learned models can accurately predict the different stay probabilities (Figure 6E, dotted lines) and the average transition speeds of the cells (Figure 6F, dotted lines). Interestingly, we observed a clear difference in the effective force *F*(*x*, *v*) of the two different miR-200c expression states of the MDA-MB 231 Tripz 200c cells which allows us to explain the higher stay probability in the miR-200c positive MDA-MB 231 cells (Figure 6G): From a dynamical systems point of view, MDA-MB 231 cells without miR-200c expression, perform so-called “limit cycle oscillations”, which means that cells repeatedly and deterministically hop between the two islands (Figure 6H). In this case, the stochasticity only affects the hopping speed. In contrast, the miR-200c positive MDA-MB 231 cells (Tripz 200c +DOX) displayed a qualitatively different behavior. Instead of deterministic oscillations, they are bistable and thus require fluctuations to hop between the islands (Figure 6I). This gave rise to less frequent and slow random transitions as observed (Figure 6E, F). Similar behavior has also been observed in the previously analyzed non-cancerous human epithelial breast cell line MCF10A (37), which endogenously expresses high levels of miR-200c (47), indicating that induction of miR-200c in the MDA-MB-231 cells fundamentally changed the phenotypic migratory behavior in our confining microenvironment. With the induction of miR-200c the motility of the breast cancer MDA-MB 231 cells converged to the motility behavior of the non-cancerous MCF10A cells (37).

Additionally, an effect of the miR-200c-inducer DOX on confined cell motility needed to be excluded. Therefore, induced MDA-MB 231 control cells (Tripz Ctrl) were examined under similar conditions. As expected, we found that the behavior of induced control cells (Tripz Ctrl +DOX, Supplementary Figure S3A to C and Supplementary Video S5) is comparable with the behavior of miR-200c negative cells (Tripz 200c -DOX) which is well described by limit cycle oscillations and is additionally comparable with wildtype MDA-MB 231 cells (37). The analysis of the effective force *F*(*x*, *v*) of induced control cells revealed a similar pattern as the uninduced MDA-MB 231 Tripz 200c cell line (Figure 6G, Supplementary Figure S3D). Thus, the administration of DOX had in general no influence on the migratory profile of MDA-MB 231 cells. In our second cell system neither MCF7 wildtype nor miR-200c knockout (KO) cells were able to perform any form of cell hopping in this dumbbell assay (Supplementary Figure S3E and F, Supplementary Video S6 and S7) and therefore an evaluation of this cell system was not possible. Taken together, our dumbbell-micropatterns provide conceptual insight into the hopping dynamics altered due to miR-200c activation: From a dynamical systems perspective, miR-200c qualitatively changed the behavior of MDA-MB 231 from deterministic hopping to stochastic and less frequent hopping within our two-state micropatterns. This indicates that miR-200c hampers the ability of cells to efficiently migrate in confining microenvironments.

## Discussion

Metastasis formation is a complex process and is best described by the invasion-metastasis cascade which summarizes mechanisms like motility, EMT, invasion, intra- and extravasation, anoikis and metastatic colonization (9). Consequently, such a complex process is less likely to be controlled by regulation of single genes, but rather on the level of miRNAs due to the pleiotropy of their targets. Therefore, our goal was to find out whether the 23 nucleotides of miR-200c are sufficient to impede metastasis formation. In this study, we utilized a mouse xenograft model with inducible miR-200c to investigate the biological complexity of metastasis. At present, there is a lot of information published demonstrating that miR-200c expression influences different physiological parameters, like EMT, adhesion, motility etc., which all could have an impact on metastasis. However, only few studies performed *in vivo* experiments modulating the miR-200c expression. The aims of these studies focused mainly on tumor growth, chemoresistance or immune-modulatory effects (30, 32, 33, 48–52). The major metastatic sites of breast cancer are the bone, lung, brain and liver (53–56) which display significant hurdles to an effective breast cancer treatment. The reason for the limited treatment option of metastatic breast cancer is for example lung metastasis resulting in a median survival of about 25 months, only (57, 58). In our study, we analyzed these distant organs *ex vivo* for luciferase activity and thus metastasis formation. Induction of miR-200c expression in the TNBC primary tumors resulted in diminished rates of secondary tumors in all organs which we quantified as the metastatic burden, representing the number of organs infiltrated with metastases. Hence, we discovered an inhibitory effect on metastasis formation for miR-200c specifically, whereas other studies dealing with cell spreading used either the whole miR-200c/141 cluster, miR-200b or functional families including three different miRNAs (32, 59–64). According to the study of Simpson *et al.* where the miR-200c/141 cluster was analyzed, MXRA8 is a miR-200 target, at least *in silico*, and a factor in breast cancer metastasis formation (32). Pulmonary metastases in this study showed elevated expression levels of MXRA8, which is associated with poor distant metastasis free survival in patients with basal-like breast cancer. There is also some evidence that miR-200c has anti-metastatic effect in the clinics. Song *et al.* analyzed the distant metastasis in 134 breast cancer patients and revealed that it was more likely to find patients with distant metastases in the group of patients with low levels of miR-200c (49). Moreover, our lab correlated a decreased overall survival of breast cancer patients, which especially in mammary tumors is connected with metastases, with low miR-200c expression (21). Taken together we strongly propose miR-200c as a metastasis suppressor *in vivo*.

Interestingly, we found that miR-200c alone is able to hamper the complex process of metastases formation. This might be possible by directly affecting several important miR-200c targets like XIAP, FOXF2, ZEB1, E-Cadherin, HMGB1, MXRA8, ZEB2, FN1, LEPR, FHOD1, PPM1F, Moesin, NTRK2, BMI-1, KRAS, PDE7B, USP25 and further as reviewed by i.e., Humphries *et al.* (1, 28, 32, 49, 52, 61, 65–70). MiR-200c inhibits EMT and promotes MET by means of decreasing the expression of zinc finger E-box-binding homeobox 1 and 2 (ZEB1 and ZEB2) through direct binding which subsequently results in the increase of E-cadherin expression (1, 18). As a consequence, cancer cells maintain an epithelial, non-migratory behavior when miR-200c is expressed. But ZEB1 and ZEB2 themselves are able to reverse the transition of cancer cells into a mesenchymal-like, highly motile character, as both transcription factors can bind to the E-box sites of miR-200c and additionally hamper the transcription of E-cadherin (1). However, our goal was not to decipher the contribution of the many miR-200c targets to cell spreading, but to focus on cellular migration, which is the prerequisite of metastases. We reduced the complexity of the cellular systems gradually and explored whether we eventually see effects of miR-200c on single cells in a confined environment. When analyzing collective cell migration, we observed a confluence dependent effect of miR-200c on the motility. At higher confluence rates cell-cell contacts and cellular adhesion are very likely to influence the migratory speed. Only the use of novel tools to monitor undirected migration made it possible to make a valid statement on cell motility, which we determined to be dependent on the confluence of the cells. We found that in epithelial MCF7 cells the enhanced speed of miR-200c KO cells is reduced most probably due to cell-cell contacts and cellular adhesion. Moreover, the growing behavior in large clusters underlines the importance of cell-cell interactions for epithelial cells. In the mesenchymal MDA-MB 231 cells the motility is even increased when miR-200c is expressed and cells become denser. We hypothesize that upon miR-200c expression cell surface molecules like E-Cadherin (71) are expressed and facilitate an improved sliding of cells instead of uncontrolled cell collisions when miR-200c is absent. Cell-cell interactions could also explain why single cells on the dumbbell-shaped geometry do not show such an increase in motility. Thus, the cellular phenotype (epithelial or mesenchymal) seems to be of great importance for the collective migration of cells. Shown by Kopp *et al.* miR-200c expression is tightly corresponding to the phenotype of breast cancer cells. While mesenchymal breast cancer like MDA-MB 231 cells show no or low expression of miR-200c, epithelial cells as MCF7 are characterized by a high expression of miR-200c (26). With the utilization of different cellular models like a stable knockout or inducible overexpression of miR-200c in breast cancer cells, phenotypes were switched within a single cell line and allowed us a comprehensive analysis of collective and single cell migration (22, 23).

We sought to acquire deeper insights into this mechanism by using scratch assays, where cells could move unrestricted in the free area of the scratch. Wound closure as well as the number of invaded cells was enhanced in miR-200c non-expressing breast cancer cells. The analysis of the confinement ratio (also known as straightness index or persistence (72, 73)) and directionality, which indicate how “efficient” a cell is moving from its starting point in a direction, revealed miR-200c negative cells to show increased directed movements (74). Especially when the motility behavior of cells in the front and bulk region of the scratch were analyzed, miR-200c negative cells at the front showed elevated migration towards the free space in the scratch. MiR-200c positive cells however tend to stay in their cellular clusters. This is particularly important as we observed no significant difference in migration speed. Here again we ascribe the altered migratory behavior to cell contacts, which seem to have a great impact on the direction of migration. An epithelial monolayer, as mimicked by miR-200c expressing MDA-MB 231 cells, usually moves in a coordinated way while mesenchymal cells show more individual collective behavior (75). As this is not the case in miR-200c expressing cells, this miRNA hampers cell motility but does not induce the ability to form collectively migrating monolayers.

To investigate the transition of collective cell migration to single cell movement we used transwell assays. Here a cell needs to break out of its cluster to cross a membrane through a pore. By doing so it additionally needs to modify its shape, demonstrating cellular plasticity. Our findings show a hampered migratory behavior of miR-200c positive cells through the membrane of the transwell. These results were supported by Jurmeister *et al.* who showed a decrease in invasion when MDA-MB 231 cells were transfected with miR-200c mimics in such a setting (69). A significant reduction as observed in our study was also reported by Park *et al.* though MDA-MB 231 cells being transfected with miR-200a/c (76).

Finally, a key aspect of metastasis is how single cells navigate complex confining geometries, when they migrate away from the initial tumor site (17). Therefore, we studied cell migration in a simple 1D assay that confined cells to frequently squeeze through a thin constricting bridge. In this system, we observed that miR-200c negative cancer cells perform deterministic transitions between the islands, while miR-200c positive cancer cells only transition randomly. These dynamics result in less frequent and slower transitions for miR-200c positive cancer cells, indicating that miR-200c is a key player of regulating confined cell migration and invasiveness of cells. A key aspect of cell migration of the wildtype MDA-MB 231 on the dumbbell-shaped geometry is that the bridge acts as a cue for the cell to continuously squeeze through (37). Flommersfeld *et al.* and Brückner *et.al*. (77, 78) suggest that this might be due to increased polymerization of actin fibers at the cell front, when the cell’s protrusion is inside the bridge. This might be due to increased alignment of actin fibers (79) or altered diffusion of polarity cues (80) when the protrusion is laterally confined. Thus, we hypothesize that changes in the cytoskeleton by miR-200c expression lead to less cellular protrusions on the bridge and a decreased intracellular force on the nucleus to follow these protrusions onto the bridge. One key player that could indirectly be repressed by miR-200c is a crosslinker of the actin cytoskeleton Filamin A. Lack of this crosslinker could lead to morphological changes and reduced cellular motility (22). MiR-200c can also alter the expression of vimentin which is an intermediate filament (IF) protein (81). The absence of miR-200c increases vimentin expression (82, 83) which in turn is recently reported to promote cell migration (81). Given the multitude of potential targets of miR-200c, further research is needed to disentangle the role of miR-200c in confined cell migration. However, our experiment clearly shows that miR-200c negatively affects the ability of cancer cells to efficiently squeeze through constricting obstacles.

The epithelial MCF7 cell line did not show hopping on the dumbbells indicating less cytoskeletal dynamics like protrusions etc. Mathieu *et al.* analyzed the motility behavior of MCF7 cells on microlines and discovered only small velocity values (84) which can be interpreted as nearly no active movement of single MCF7 cells. The hurdle of crossing the bridge of the dumbbell assay in our case might be even more insuperable for these non-metastatic cells. Nevertheless, both single cell motility assays, namely the transwell and the dumbbell assay, are showing decreased migratory behavior of miR-200c positive cells.

## Conclusion

In conclusion, this study aimed to connect the observed reduction of the metastatic burden due to miR-200c *in vivo* to effects of miR-200c on cell migration *in vitro*. To this end, we reduced the complexity of metastasis to a series of established and novel model systems for cell migration. Importantly, the different migration assays vary in scale ranging from collective cell migration to single cell migration in structured confining microenvironments. While a complete picture of the effect of miR-200c remains to be unraveled, our experiments give insight into the physical role of miR-200c on key aspects of cell migration. In particular, we observed in all experiments a reduction of cell motility that manifested in reduced wound healing, less frequent dissociation of single cells from a cluster through pores, and less frequent and slower transition dynamics of confined cell migration. Of note, as miR-200c is sufficient to prevent metastasis formation it possesses the right range of pro-metastatic targets and hence can be termed a metastasis suppressor. Together with a more mechanistic understanding of how miR-200c affects the migratory machinery, understanding cell-cell interactions could give deeper insight into how miR-200c regulates cell migration on an individual and collective scale. Our study provides a systematic, conceptual and quantitative characterization of how intracellular components of cells determine key aspects of metastasis. Future work will show whether the administration of miR-200c nucleic acid therapeutics might provide an innovative way to prevent metastases in breast cancer.

## Acknowledgments

The authors want to thank Enikö-Melinda Kiss and Lorina Bawej for the support in cell culture work, Jonathan García-Roman for performing the TALENs knockout, and Konstantin Schaffer from PHIO scientific GmbH for his support using their technology. Furthermore, the authors want to thank David B. Brückner and Johannes Flommersfeld for helpful discussions.

## Video Description

Video 1: Undirected migration of MCF7 wt cells. Live cell imaging was performed with the Cellwatcher M.

Video 2: Undirected migration of MCF7 KO 200c cells. Live cell imaging was performed with the Cellwatcher M.

Video 3: Undirected migration of MDA-MB 231 Tripz 200c -DOX cells. Live cell imaging was performed with the Cellwatcher M.

Video 4: Undirected migration of MDA-MB 231 Tripz 200c +DOX cells. Live cell imaging was performed with the Cellwatcher M.

Video 5: Wound closure of MDA-MB 231 wt stained with 1 µM siR-DNA. The timeframe from 0 to 10 hours was monitored.

Video 6: Trajectories of migrating MDA-MB 231 wt cells. The timeframe from 0 to 10 hours was monitored.

Video 7: Wound closure of MDA-MB 231 Tripz 200c +DOX stained with 1 µM siR-DNA. The timeframe from 0 to 10 hours was monitored.

Video 8: Trajectories of migrating MDA-MB 231 Tripz 200c +DOX cells. The timeframe from 0 to 10 hours was monitored.

Video 9: MDA-MB 231 Tripz 200c (-DOX) moving on the two-state dumbbell micropattern coated with fibronectin. Nucleus is stained in blue.

Video 10: MDA-MB 231 Tripz 200c (+DOX) moving on the two-state dumbbell micropattern coated with fibronectin. Nucleus is stained in blue. Induced miR-200c and RFP expression shown in red.

## Supplementary Videos

Video S1: Wound closure of MCF7 wt stained with 1 µM siR-DNA. The timeframe from 14 to 24 hours after scratch induction was monitored.

Video S2: Trajectories of migrating MCF7 wt cells. The timeframe from 14 to 24 hours was monitored.

Video S3: Wound closure of MCF7 KO 200c stained with 1 µM siR-DNA. The timeframe from 14 to 24 hours after scratch induction was monitored.

Video S4: Trajectories of migrating MCF7 KO 200c cells. The timeframe from 14 to 24 hours was monitored.

Video S5: MDA-MB 231 Tripz Ctrl (+DOX) moving on the two-state dumbbell micropattern coated with fibronectin. Nucleus is stained in blue. Induced scrambled control sequence and RFP expression shown in red.

Video S6: MCF7 wt moving on the two-state dumbbell micropattern coated with fibronectin. Nucleus is stained in blue.

Video S7: MCF7 KO 200c moving on the two-state dumbbell micropattern coated with fibronectin. Nucleus is stained in blue.

## Author Contributions

Conceptualization, B.K. and A.R.; methodology, B.K., E.B., T.B., E.H., U.W., J.P., A.J., P.P., S.Z., E.W. and A.R.; software, E.B., T.B.; validation, B.K., E.B., T.B., U.W., J.P., A.J., S.Z. and A.R.; formal analysis, B.K., T.B.; investigation, B.K., E.B., T.B., U.W., J.P. and E.W.; resources, B.K., E.B., T.B., U.W., J.P. and S.Z.; data curation, B.K., E.B., T.B., A.J. and A.R.; writing—original draft preparation, B.K., E.B., T.B. and A.R.; writing—review and editing, B.K. and A.R.; visualization, B.K. and A.R.; supervision, P.P., S.Z., C.P.B., J.O.R., E.W. and A.R.; project administration, A.R.; funding acquisition, E.W. and A.R. All authors have read and agreed to the published version of the manuscript.

## Institutional Review Board Statement

All animal experiments were performed according to the guidelines of German law for the protection of animal life and were approved by the district government of Upper Bavaria. Reference number: ROB-55_2-2532_Vet_02-19-20.

## Data Availability Statement

All stable cell lines generated in this study are available from the corresponding author upon reasonable request. The data generated in this study are available on request from the corresponding author.

