## Supplementary figures and figure legends for "Unravelling the metastasis-preventing effect of miR-200c *in vitro* and *in vivo*"

### Supplementary Figure S1

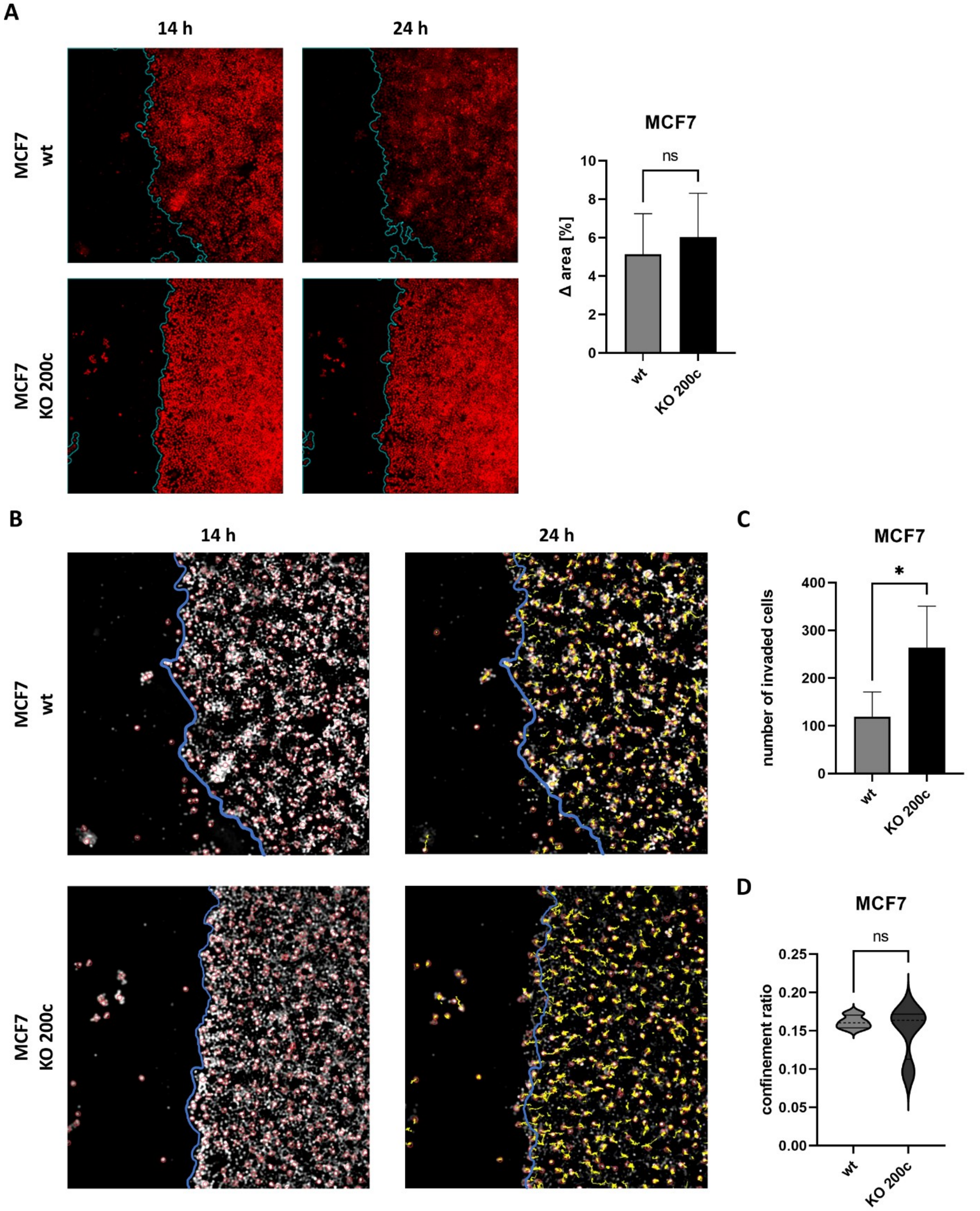


**Supplementary Figure 1: MCF7 breast cancer cells lacking miR-200c expression tend to leave cell clusters more frequently. (A)** Wound closure of MCF7 cells with (wt) or without (KO 200c) miR-200c expression. Blue lines in the microscopic pictures (left) indicate the borders of the wound. MCF7 cells tended to get stacked on top of each other when performing the scratch and later on were rolling out slowly. Therefore, the wound closure was monitored for 10 hours but starting 14 hours post scratching. The difference in the area in [%] (right) is used to quantify the scratch closure (n = 4). Cells are stained with siR-DNA. **(B)** Microscopic pictures of MCF7 cells with different miR-200c expression for the analysis of **(C)** the number of invaded cells and **(D)** the confinement ratio (n = 4). Blue lines represent the border of the scratch at 14 hours. Yellow lines in the microscopic pictures represent the trajectories of the cells at 24 hours after scratch. The dashed line shows the median and the full lines the quartiles of the violin plot in (D). For statistical evaluation an unpaired, two-tailed student's t-test was performed. ns = not significant, * $p < 0.05$.

### Supplementary Figure S2

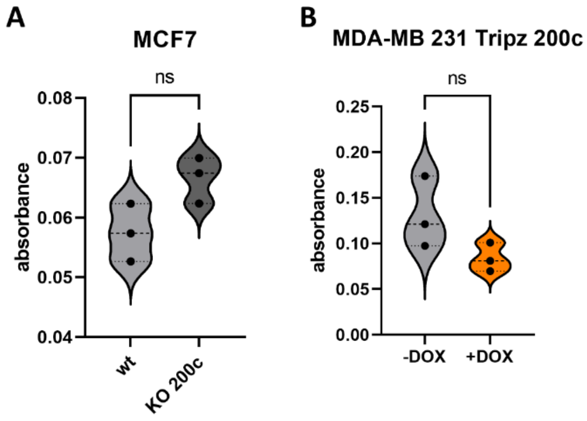


**Supplementary Figure S2: Absorbance measurement of crystal violet stained, migrated cells reveals enhanced migration of miR-200c negative cells.** The absorbance of crystal violet stained **(A)** MCF7 (n = 3) and **(B)** MDA-MB 231 Tripz 200c cells (n = 3) was measured. The amount of crystal violet incorporated by the individual migrated cells was determined and finally displayed as a violin plot with individual values. For statistical evaluation an unpaired, two-tailed student's t-test was performed. ns = not significant

### Supplementary Figure S3

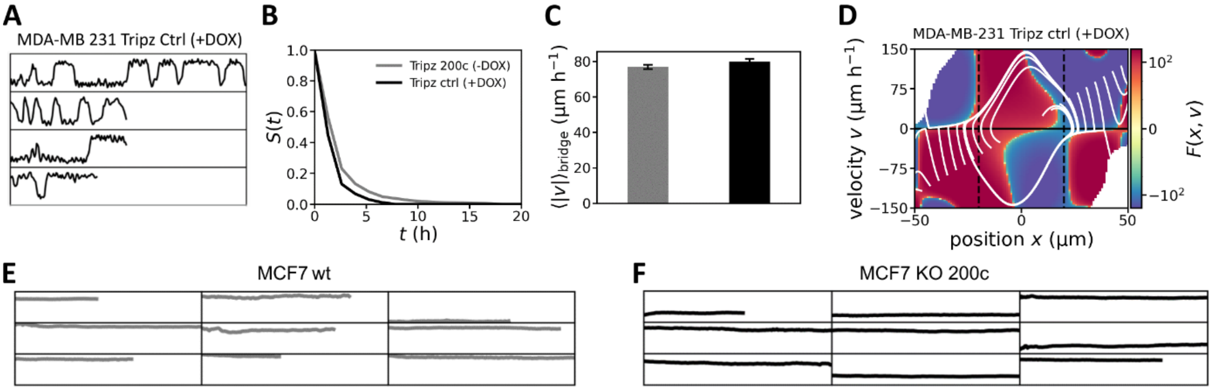


**Supplementary Figure S3: Doxycycline induction is not affecting confined cell migration of mesenchymal cells. No measurable hopping dynamics in epithelial MCF7 cells.** Selection of nucleus trajectories for **(A)** MDA-MB 231 Tripz Ctrl +DOX cells. **(B)** Stay probability of the miR-200c non-expressing cell lines. Comparison of miR-200c negative cell lines (MDA-MB 231 Tripz Ctrl +DOX, black and MDA-MB 231 Tripz 200c -DOX, gray). **(C)** Average speed of the MDA-MB-231 Tripz Ctrl (+DOX) and MDA-MB-231 200c (-DOX) cells while making a transition on the bridge. **(D)** Inferred deterministic part of the hopping dynamics. White lines show the deterministic behavior in the two-dimensional phase space of the cells for MDA-MB 231 Tripz Ctrl +DOX. Selection of nucleus trajectories for **(E)** MCF7 wt and **(F)** MCF7 KO 200c cells.
