## Supplementary methods for "Unravelling the metastasis-preventing effect of miR-200c *in vitro* and *in vivo*"

**Supplementary Material and Methods**

1. Single cell motility on dumbbells

For experiments with MCF7 10,000 cells were seeded on the micropatterned ibiTreat µ-dish (ibidi GmbH) and left to adhere for at least 4 hours. The further implementation of the experiments corresponds to the method already mentioned in the main part besides the size of the micropattern for MCF7 which was adjusted and 1.3x bigger than that of MDA-MB 231 cells. This was because MCF7 cells are approximately 1.3x bigger than MDA-MB 231 cells. 62 tracks of the MCF7 wt and 44 tracks of the MCF7 KO 200c cells were analyzed. Each single measurement was conducted for at least 48 hours, but track length varied between 8 hours (minimum) and 48 hours (maximum). Every 10 minutes one image was taken.
